# Patients with ventromedial prefrontal lesions show an implicit approach bias to angry faces

**DOI:** 10.1101/2020.06.21.162628

**Authors:** Macià Buades-Rotger, Anne-Kristin Solbakk, Matthias Liebrand, Tor Endestad, Ingrid Funderud, Paul Siegwardt, Dorien Enter, Karin Roelofs, Ulrike M. Krämer

**Author notes:** Corresponding author: Macià Buades-Rotger, PhD, Radboud University, Montessorilaan 3, 6525 HR Nijmegen, The Netherlands.

## Abstract

Damage to the ventromedial prefrontal cortex (VMPFC) can cause maladaptive social behavior, but the cognitive processes underlying these behavioral changes are still uncertain. Here, we tested whether patients with acquired VMPFC lesions show altered approach-avoidance tendencies to emotional facial expressions. Thirteen patients with focal VMPFC lesions and 31 age- and gender-matched healthy controls performed an implicit approach-avoidance task in which they either pushed or pulled a joystick depending on stimulus color. While controls avoided angry faces, VMPFC patients displayed an incongruent response pattern characterized by both increased approach and reduced avoidance of angry facial expressions. The approach bias was stronger in patients with higher self-reported impulsivity and disinhibition, and in those with larger lesions. We further used linear ballistic accumulator modelling to investigate latent parameters underlying approach-avoidance decisions. Controls displayed negative drift rates when approaching angry faces, whereas VMPFC lesions abolished this pattern. In addition, VMPFC patients had weaker response drifts than controls during avoidance. Finally, patients showed reduced drift rate variability and shorter non-decision times, indicating impulsive and rigid decision-making. Our findings thus suggest that VMPFC damage alters the pace of evidence accumulation in response to social signals, eliminating a default, protective avoidant bias and facilitating dysfunctional approach behavior.

## Introduction

Patients with damage to the ventromedial prefrontal cortex (VMPFC) often show disruptive social behavior (Anderson et al., 2006; Barrash et al., 2000; Beer et al., 2006; Blair, 2004). Ventromedial lesions typically impact adjacent white matter, thereby hindering VMPFC-amygdala cross-talk (Folloni et al., 2019) and rendering individuals more emotionally reactive (Jenkins et al., 2018; Motzkin et al., 2015). Consequently, antisocial behavior related to VMPFC dysfunction has been classically attributed to deficits in emotion regulation (Davidson et al., 2000). However, this view has proven difficult to reconcile with the many other functions ascribed to the VMPFC, such as subjective value computation (Clithero & Rangel, 2014). This apparent discrepancy has been mended by recent theories that conceptualize emotion regulation as a special case of value-based decision-making, wherein the brain must choose among mutually contradicting affective states and behaviors (Dixon et al., 2017; Gross, 2015; Koch et al., 2018). Recent investigations adhere to this model in ascribing an *evaluative* and *generative* role to the VMPFC (Hiser & Koenigs, 2018). According to this view, the VMPFC codes for the potential hedonic or threatening value of a given stimulus in order to steer the organism towards or away from it (Rudebeck & Rich, 2018). In this framework the VMPFC is assumed to generate cognitive maps of current internal states and external sensory information, enabling the selection of the most appropriate course of action (Stalnaker et al., 2015; R. C. Wilson et al., 2014). Such a process has been termed model-based or goal-directed behavior because it operates on the basis of internal representations of oneself and the environment rather than by force of habit (Lucantonio et al., 2012).

From this rationale, it follows that antisocial behavior after VMPFC damage could arise from inaccurate assessment and selection processes. More specifically, VMPFC lesions might impair the ability to correctly predict the consequences of one’s own actions in response to social signals (Rudebeck & Murray, 2014), e.g., wrongly expecting rewards from approaching potential punishment cues. Nevertheless, evidence to support this tenet is scarce in humans with VMPFC lesions. One report suggests that VMPFC-damaged patients display an altered sense of personal distance, e.g., they get closer to strangers (Perry et al., 2016). Comparably, a study showed that persons with VMPFC lesions judge negative facial expressions (i.e., angry, disgusted, fearful and sad) as *more* approachable (Willis et al., 2010). It remains to be tested, however, whether these tendencies can be attributed to implicit biases during action selection, and whether these putative alterations are linked with actual impairments in daily functioning. Moreover, it is unclear which precise cognitive mechanisms underlie such abnormal behavioral dispositions. These are important steps in understanding how VMPFC-dependent disturbances in social behavior play out in everyday life.

In order to clarify these issues, we investigated whether VMPFC lesions lead to implicit response biases towards or away from negative, positive, or neutral facial expressions. We used a version of the approach-avoidance task (AAT) wherein subjects have to either push or pull a joystick depending on the color (e.g. red or green) of a human face (Roelofs et al., 2010). Faces are programmed to grow or shrink in size accordingly, giving the impression that they loom closer or recede upon pulling and pushing, respectively. Hence, the AAT allows measuring implicit response tendencies to task-irrelevant features of the faces such as their emotional expression. A study with this task suggested that psychopaths lack automatic avoidance of directly-gazing angry faces, and that this effect was correlated with aggressiveness (von Borries et al., 2012). Following a similar approach, we tested whether task scores correlated with patients’ daily emotional behavior as measured with validated clinical scales in order to assess the practical relevance of possible approach-avoidance biases.

In addition, we scrutinized the putative cognitive mechanisms underlying altered task performance in VMPFC patients using Linear Ballistic Accumulator (LBA) modelling on response times (Brown & Heathcote, 2008). LBA modelling assumes that decisions arise from a sequential evidence accumulation process, the speed of which is determined by multiple latent variables (e.g., pre-existing response tendencies or shorter decision latencies) that can be quantified and compared between experimental conditions and/or groups. Previous modelling studies on an explicit version of the AAT reported relatively faster evidence accumulation in healthy subjects when threatening stimuli are to be avoided (Krypotos et al., 2015; Tipples, 2019). LBA modelling might hence offer insights not captured by standard methods of reaction time analysis.

## Materials and methods

### Participants and lesion localization

The clinical sample consisted of 13 patients with chronic (> 6 months post-injury or surgery), focal damage to the ventral prefrontal cortex (mean age=50.8 [27-62], 7 women, 12 right-handed). Lesions were mainly ventromedial (Fig. 1A; Table 1), namely Brodmann Areas (BA) 10, 11, 24, 25, and 32. There was also substantial rostromedial damage (BA 9 and anterior BA 10) and, to a lesser extent, in the ventrolateral prefrontal cortex (BA 45, 46, and 47). Harm to posterior dorsomedial areas (BA 6 and 8) was minimal, with a small number of lesions extending to the anterior insula (BA 13). Lesions were predominantly right-sided in ten patients, one had an exclusively left-sided lesion, and two had comparable damage in both hemispheres. All lesioned tissue was restricted to the frontal lobe. Etiology of the lesions was either meningioma (n=9), traumatic brain injury (n=2), oligodendroglioma (n=1), or astrocytoma (n=1). The control sample was composed of 31 age- and gender-matched neurologically healthy individuals (mean age=50.1 [43-54], 19 women, all right-handed). As previously reported, patients had normal or corrected to normal vision, showed no deficits in standard neuropsychological testing, and had no motor dysfunction of the hands. However, they reported greater difficulties in executive function, metacognition, and behavioral regulation as compared to a separate control sample (see Løvstad et al., 2012 for a complete report). All patients were recruited and measured at Oslo University Hospital and the University of Oslo, whereas the behavioral control sample was recruited and measured at the University of Lübeck. All participants provided informed consent and the study procedures adhered to the Declaration of Helsinki. The study was approved by the ethics committee of the University of Lübeck and the Regional Committee for Medical Research Ethics - South East Norway.

**Figure 1.**
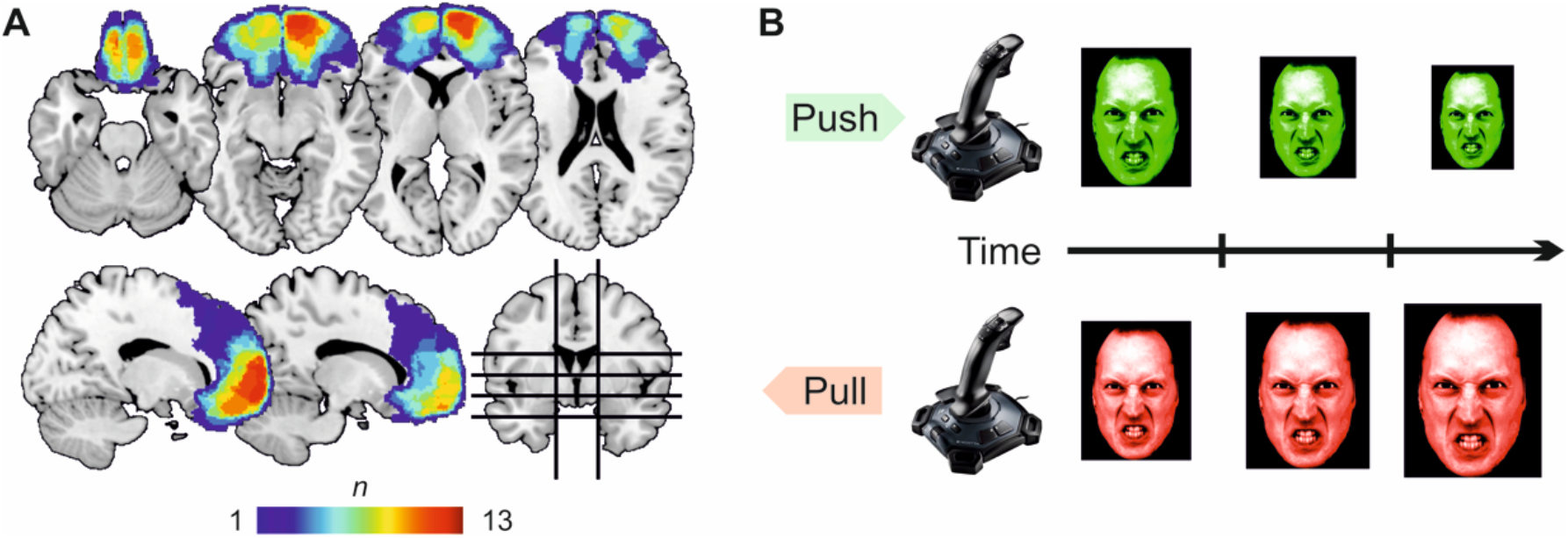
**A)** Lesion overlap. Warmer colors depict more overlap between patients. Peak overlap was located in x=4, y=58, z=−14 (Montreal Neurological Institute space). **B)** Schematic depiction of the implicit Approach Avoidance Task (AAT). Subjects had to either push or pull a joystick in response to the color of the presented face while ignoring its facial expression (angry, happy, or neutral), gaze (direct or averted), gender (male or female), and identity (eight actors). Pushing made faces shrink in size, whereas pulling made them grow larger. The 384 trials were self-paced.

**Table 1:** Lesioned Brodmann areas

| Brodmann area | N | % damage ($M \pm SE$) |
| --- | --- | --- |
| 6 | 3 | $0.99 \pm 0.89$ % |
| 8 | 3 | $4.87 \pm 4.30$ % |
| 9 | 12 | $5.07 \pm 1.93$ % |
| 10 | 12 | $18.44 \pm 3.35$ % |
| 11 | 10 | $7.62 \pm 2.16$ % |
| 13 | 4 | $0.55 \pm 0.46$ % |
| 24 | 9 | $3.72 \pm 0.90$ % |
| 25 | 8 | $18.73 \pm 3.54$ % |
| 32 | 12 | $20.97 \pm 4.48$ % |
| 45 | 4 | $0.85 \pm 0.44$ % |
| 46 | 6 | $2.97 \pm 1.79$ % |
| 47 | 7 | $1.95 \pm 0.95$ % |

N patients: number of patients (out of 13) with damage in a given region; % damage: average percentage of lesioned tissue across patients; M: mean; SE: standard error.

### Clinical scales

Patients filled out the self-report form of the Behavior Rating Inventory of Executive Function – Adult version (BRIEF-A; Roth et al., 2005) and the Urgency, Premeditation, Perseverance, Sensation Seeking (UPPS) Impulsive Behavior Scale (Whiteside & Lynam, 2001), both ad-hoc translated into Norwegian. The BRIEF-A is a standardized rating scale consisting of 75 items that tap into everyday executive functioning within the past 6 months. Internal consistency and test-retest reliability of the BRIEF-A are reportedly high and construct validity has been established in healthy and clinical populations (Waid-Ebbs et al., 2012). For the purposes of this study we discarded all BRIEF-A scales not directly related to the control of automatic emotional tendencies (“Working Memory”, “Plan/Organize”, “Organization of Materials”, “Shift” and “Initiate”) and thus only considered the scales “Inhibit”, “Emotional Control”, and “Self-Monitor”. The Inhibit scale measures deficits in inhibitory control and impulsivity; the Emotional Control scale assesses a person’s inability to regulate emotional responses; and the Self-Monitor scale evaluates difficulties in social or interpersonal awareness. The UPPS Impulsive Behavior Scale (Whiteside & Lynam, 2001) is a 45-item self-report, assessing different facets of impulsivity on four subscales. The UPPS has been shown to display good internal consistency and construct validity (Whiteside et al., 2005). We used the total UPPS score for correlational analyses because we deemed all subscales (“Negative urgency”, “Lack of Premeditation”, “Lack of Perseverance”, ”Sensation Seeking”) to be theoretically associated with emotional action control.

### Implicit Approach-Avoidance Task (AAT)

Subjects performed the implicit approach-avoidance task (AAT; Fig. 1B) as previously described (Roelofs et al., 2010; von Borries et al., 2012). Stimuli were photographs (Ekman & Friesen, 1976; Lundqvist et al., 1998) showing the face of one out of eight actors (four male and four female) displaying angry, happy or neutral expressions with either direct (straight) or averted (sideways) gaze. Photographs were cut out ovally and tinted red or green, amounting to a total number of 384 trials. Participants performed 18 practice trials comprising only straight-gazing neutral faces, followed by the experimental trials. After half of the trials, subjects had a break, performed two additional practice trials with directly gazing neutral expressions to recall task demands and completed the second half. Stimuli were presented randomly, with no more than three of the same emotion-response combinations in succession. The neutral faces from the practice trials were also presented in the task proper.

Pictures were presented at a 1024 x 768 pixels resolution on a computer screen. We placed the joystick (Logitech Attack 3) between subject and screen to allow for comfortable pull and push movements. Participants started each trial by pressing the fire button with the index finger of the dominant hand. A face stimulus appeared in the center of the screen. Participants were instructed to ignore the facial expression and only respond to the color of the face. Half the participants had to push the joystick in response to red and pull in response to green stimuli, the other half had the opposite instruction. To visually emphasize that pull movements meant approach, and push movements meant avoidance, pictures grew or shrank in size following pull or push movements, respectively. Stimuli had a starting size of 9.5° by 13° and could shrink to a minimum of 3.5° by 4.5° when pushing or grow to a maximum of 15.5° by 20° when pulling. In practice trials, pictures remained visible after erroneous responses to allow for response correction, whereas in the task proper stimuli disappeared after they had reached minimal or maximal size. Participants were instructed to respond as quickly and accurately as possible. Importantly, trials could only be initiated once the joystick was placed back in its original centered position.

### Behavioral data analysis

Reaction times (RT) were recorded as time from stimulus onset until the first joystick movement. We excluded incorrect trials as well as those with RT shorter than 150ms or longer than 1000ms, and extracted mean log-transformed RT per cell (as in Bertsch et al., 2018). We then ran an analysis of variance (ANOVA) on the resulting values with within-subject factors emotion (happy, neutral, angry), actor gender (male, female), gaze (left, right, and direct), movement (pull or push), and the between-subject factor group (VMPFC vs healthy controls) using the *ez* package (version 4.4-0). We modelled all relevant task factors as in previous studies with the implicit AAT (Roelofs et al., 2010; von Borries et al., 2012). In order to control for multiple testing, we applied a False Discovery Rate (FDR) correction as recommended for exploratory ANOVAs (Cramer et al., 2016). Color and condition were counterbalanced across participants (green=pull for one half, green=push for the other half) and are thus controlled for by design, though this randomization was not stratified by gender or other participant characteristics. We inspected significant effects with post-hoc t-tests.

Error rates in the implicit AAT are often low, due to which between-condition differences in error rates are not analyzed (Roelofs et al., 2010; von Borries et al., 2012). Here, too, errors were few and unevenly distributed across conditions. Therefore, as in previous studies (Roelofs et al., 2010; von Borries et al., 2012), we simply compared the mean error rate between groups. We used Welch’s t-test, which is robust to unequal variances and uneven sample sizes (Ruxton, 2006). Subsequently, we computed Pearson correlation coefficients between AAT scores (between-condition differences in RT and overall error rates) and each of the four clinical scales. We assessed the robustness of significant correlations with bootstrap resampling to obtain 95% bias-corrected accelerated confidence intervals (BCa CI) with 10000 iterations using the *bootstrap* package (version 2019.5). We performed all analyses described in this section in R (version 3.6.1) running on R Studio (version 1.1.423).

### Linear ballistic accumulator (LBA) modelling of reaction times

We subsequently implemented Linear Ballistic Accumulator (LBA) modelling on reaction time data (Brown & Heathcote, 2008). LBA models assume that decisions stem from a sequential evidence accumulation process (Fig. 3A). Evidence for each response option is gathered linearly by a separate accumulator, which races against the other/s until one of them reaches a decision threshold. Evidence accumulation starts after a variable period of non-decision time and its speed is given by the drift rate, which is sampled from a normal distribution. The standard deviation of this distribution constitutes what we here label drift noise, i.e., variability in the pace of evidence accumulation. In addition, the accumulators might begin each trial from a different starting point, which is drawn from a uniform distribution. Therefore, a response option will be taken more quickly if starting point and decision threshold are nearer, if the drift rate is higher and less variable, and if the non-decision time is shorter. LBA models are akin to the now-popular drift diffusion models (DDM), but are simpler and more tractable computationally and thus well-suited for the relatively low amount of trials available in the present dataset (see Heathcote and Hayes, 2012, for a detailed empirical comparison between LBA and DDM).

Here, we fitted a series of LBA models with two accumulators (approach and avoidance) and four parameters: decision threshold, starting point, drift rate, and drift noise. We tested a total of 16 models in which a given combination of these parameters was allowed to vary between the six experimental conditions of interest: pull angry, pull happy, pull neutral, push angry, push happy, and push neutral. We could not test for a modulation of experimental condition on non-decision time because models including this effect failed to converge in most subjects. See Table 2 for a summary of all models. We fitted each model on the reaction time data of each individual participant using full information maximum likelihood estimation as implemented in the *glba* package version 0.2 (https://CRAN.R-project.org/package=glba). We used raw RT excluding errors and responses quicker than 150ms or slower than 1s. For model comparison we inspected which model yielded the lowest Bayesian Information Criterion (BIC) values across participants. BIC is a standard fit measure that penalizes model complexity (Burnham & Anderson, 2004; Raftery, 1995). Our model fitting and comparison approach is highly comparable to that of a recent DDM study on social approach-avoidance decisions (Mennella et al., 2020). Afterwards, we simulated data per group using the *rlba()* function and the average parameter estimates from the winning model. We next ran an ANOVA on the parameters of the winning model with factors emotion, movement, and group in order to test which model parameters varied as a function of group (Mennella et al., 2020). Finally, we compared parameters between groups with independent-samples Welch t-tests. We used R (version 3.6.1) running on R Studio (version 1.1.423) for all analyses in this section.

**Table 2:** Summary of Linear Ballistic Accumulator models tested

| **Modulated parameters in model** | **K** | **Median BIC** |
| --- | --- | --- |
| Threshold, starting point, drift rate, drift noise | 25 | -546.16 |
| Threshold, starting point, drift rate | 20 | -575.52 |
| Threshold, starting point, drift noise | 20 | -75.84 |
| Threshold, drift rate, drift noise | 20 | -606.32 |
| Starting point, drift rate, drift noise | 20 | -606.47 |
| Threshold, starting point | 15 | -117.53 |
| Threshold, drift rate | 15 | -629.57 |
| Threshold, drift noise | 15 | -119.90 |
| Starting point, drift rate | 15 | -624.78 |
| Starting point, drift noise | 15 | -157.82 |
| Drift rate, drift noise | 15 | -629.92 |
| Threshold | 10 | -161.38 |
| Starting point | 10 | -107.24 |
| **Drift rate** | **10** | **-646.62** |
| Drift noise | 10 | -172.87 |
| Null model | 5 | -169.83 |

K: number of free parameters; BIC: Bayesian Information Criterion. The model marked in **bold** had the best fit to the data across participants.

### Neuroimaging data acquisition and analysis

Structural brain volumes were recorded at the Intervention center at Oslo University hospital (Norway) on a Philips Ingenia 3-T scanner. We acquired structural images with a T1-weighted 3D turbo gradient-echo sequence with the following settings: repetition time (TR)=1.900ms, echo time (TE)=2.23ms, flip angle=8°, voxel size=1mm^3^, field-of-view (FOV)=256×256mm. Members of the team at the University of Oslo, trained in lesion reconstruction, manually delineated lesion masks on each patient’s anatomical images. We normalized these masks as recommended for lesioned brains (Ripollés et al., 2012) and created lesion overlap maps using MRIcron (Rorden & Brett, 2000). We further inquired which Brodmann Areas (BA) were damaged, following previous work (Jenkins et al., 2014). We extracted all BA masks from the Wake Forest University PickAtlas (Maldjian et al., 2003) and co-registered them with the Montreal Neurological Institute-152 template to ensure that lesions and masks were in the same space (Jenkins et al., 2014). We then computed the number of overlapping voxels between each lesion and BA mask. For each non-intact BA, we report the number of patients with damage in that region along with the mean percentage of damaged tissue (Table 1; see also *Participants and lesion localization* above).

We also inspected whether lesion size was linked with reaction times and error rates in the task. We correlated lesion size with behavioral parameters showing a group difference in the AAT and obtained the 95% bootstrapped CIs with 10000 iterations using the *bootstrap* R package to assess these effects’ robustness. We proceeded identically with all lesioned Brodmann areas, and compared the lesion-behavior correlations with each other using the William’s test as implemented in the *r.test()* function from the *psych* R package.

## Results

### Approach-Avoidance Task (AAT) results

In our primary analysis of reaction times we observed main effects of group (F_1,42_=11.92, p=.001, pFDR=.010) and emotion (F2,84=6.89, p=.001, pFDR=.010) which were qualified by an emotion x movement interaction that did not survive multiple comparison correction (F2,84=4.36, p=.015, pFDR=.083), and, crucially, by a group x emotion x movement interaction (F2,84=12.64, p<.001, pFDR<.001). In order to dissect the latter three-way interaction, we computed the difference between push and pull (i.e., avoidance minus approach) for each emotion and inspected for differences between emotion categories in each group, following previous work (Roelofs et al., 2010; von Borries et al., 2012). As shown in Fig. 2A, VMPFC patients showed a stronger approach bias toward angry relative to both happy (t_12_=3.17, p=.008) and neutral faces (t_12_=4.32, p<.001), with no difference between happy and neutral faces (p=.416). In comparison (Fig. 2B), controls showed a trend-level avoidant bias for angry relative to neutral faces (t_30_=1.75, p=.089), with no further differences between categories (all p>.272). Thus, VMPFC patients were generally slower when pushing angry faces away relative to pulling them close.

**Figure 2.**
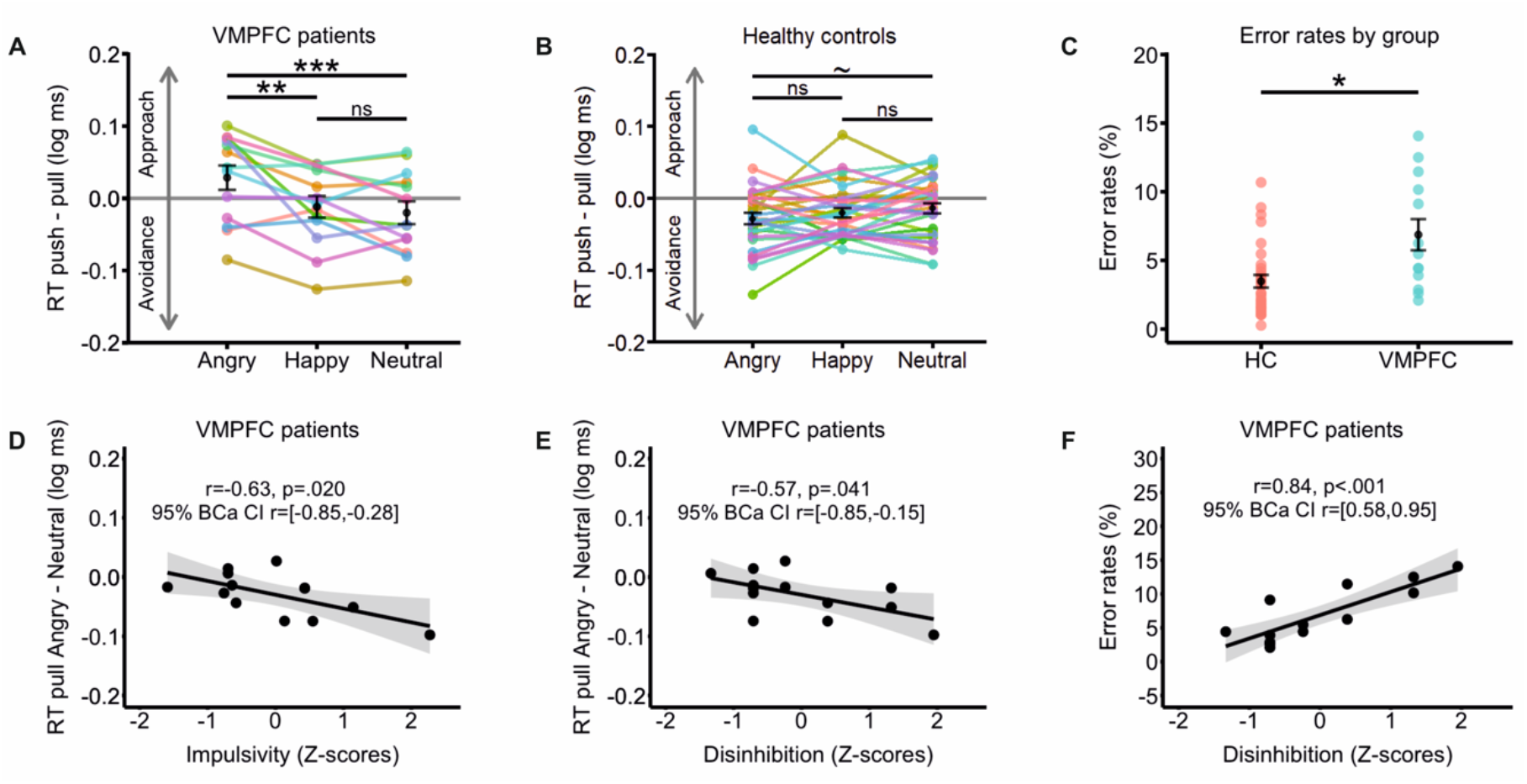
**A)** Patients with ventromedial prefrontal cortex (VMPFC) lesions showed an approach bias (reaction times [RT] for push minus pull) towards angry relative to happy and neutral faces. **B)** Healthy controls (HC) showed no bias in either direction, with a trend towards avoidance of angry relative to neutral faces. **C)** VMPFC patients made more errors than HC. **D)** Shorter RT for pull angry minus pull neutral trials were linked with greater self-rated impulsivity in VMPFC patients. **D)** Shorter RT for pull angry minus pull neutral trials were correlated with greater disinhibition in VMPFC patients. **F)** Error rates were correlated with all clinical self-reports in VMPFC patients, including greater self-rated disinhibition. ~p<.1, *p<.05, **p<.01, ***p<.001. BCa CI: Bias-Corrected accelerated Confidence Intervals obtained with bootstrapping.

In order to ascertain whether these effects were predominantly driven by approach or avoidance, we computed the difference in reaction times between emotions separately for push and pull movements in each group. Regarding approach movements, VMPFC patients were faster to pull angry relative to neutral (t_12_=3.66, p=.003) but not happy faces (p=.107). Controls showed no between-emotion differences in pull movements (all p>.278). For avoidance movements, VMPFC patients were slower to push angry relative to happy faces (t_12_=2.88, p=.013) but comparably fast when pushing angry and neutral ones (p=.284). Controls were quicker to avoid angry as compared to neutral faces (t_30_=2.27, p=.030) but not happy ones (p=.605). Therefore, controls specifically showed avoidance of angry in comparison with neutral expressions. In contrast, VMPFC patients showed increased approach of angry relative to neutral faces, and reduced avoidance of angry as compared to happy ones. We used these significant between-emotion differences for later correlation analyses, as they index the increased threat approach (pull angry minus pull neutral) and reduced threat avoidance (push angry minus push happy) demonstrated by VMPFC patients.

Additionally, there was an emotion x gaze interaction across the whole sample (F4,168=4.63, p=.001, pFDR=.011). We computed the difference in reaction times between direct and averted gaze and compared between emotions over all participants to further investigate this effect. The interaction was driven by slower reactions to directly-gazing neutral faces relative to happy (t_43_=3.09, p=.003) and, at trend level, angry ones (t_43_=1.81, p=.077).

We subsequently compared error rates between groups. Although both groups performed the task well, VMPFC patients committed about twice as many errors (6.87±1.13%) than healthy controls (3.47±0.46%), t_16.14_=2.77, p=.013 (Fig. 1C).

### Correlations between task scores and clinical scales

We then inspected for associations between clinical scales and task-derived scores, with the aim of testing the clinical relevance of approach-avoidance biases as measured with the AAT. The approach bias for angry minus neutral faces was linked with increased self-reported impulsivity (Fig. 1D; r=−.63, p=.020, 95% BCa CI=[−.85, −.28]), and greater disinhibition (Fig. 1E; r=−.57, p=.041, 95% BCa CI=[−.85, −.15]), but there were no correlations with either of the other two clinical scales, or between the angry push minus happy push difference and any of the scales (all p>.160). Error rates were associated with impairment in all scales, namely impulsivity (r=.57, p=.040, 95% BCa CI=[.05, .85]), disinhibition (Fig. 1F; r=.84, p<.001, 95% BCa CI=[.58, .95]), difficulties in emotional control (r=.59, p=.033, 95% BCa CI=[.23, .78]), and worse self-monitoring (r=.70, p=.007, 95% BCa CI=[.25, .90]).

### Correlations between lesion anatomy and AAT scores

Subsequently, we tested whether task-derived response biases were linked with lesion size. Patients with larger lesions were quicker to approach angry relative to neutral faces (r=−.77, p=.001, 95% BCa CI=[−.92, −.55]). Correlations between this bias and the percentage of lesioned tissue were negative and sizeable for each individual Brodmann area (between r=−.25 and r=−.70) and did not significantly differ from each other (all p>.099). This indicates that the tendency to approach angry as compared to neutral faces cannot be specifically attributed to any of these subregions. Lesion size was not correlated with the push angry minus push happy difference (p=.369) or with error rates (p=.469). Lesion extension was thus exclusively associated with threat approach, but not with the reduced threat avoidance and increased error rates displayed by VMPFC patients.

### Linear Ballistic Accumulator (LBA) modelling results

Next, we turned to Linear Ballistic Accumulator (LBA) modelling in order to uncover which latent decision parameters might account for VMPFC patients’ response patterns. We provide the complete list of models in Table 2. The winning model assumed that emotional expression and movement modulated drift rates exclusively. This model had the lowest Bayesian Information Criterion (BIC) across subjects (median BIC=−646.62, k=10 free parameters) and was the best-fitting model in all 13 VMPFC patients as well as in 90% (28/31) of control participants. According to model-comparison guidelines (Burnham & Anderson, 2004; Raftery, 1995), the evidence for this model can be considered substantial relative to the two next best-fitting (and slightly more complex) models, one assuming an effect of emotional expression on drift rate and drift noise (median BIC=−629.92, k=15 free parameters), and one in which emotional expression impacted drift rate and decision threshold (median BIC=−629.57, k=15 free parameters). The winning model could reproduce reaction times in pull angry trials with a precision of around ~30-50ms across successive simulations for both VMPFC patients (example mean simulated data=495ms; mean real data=544ms) and control participants (example mean simulated data=593ms; mean real data=624ms).

An ANOVA on parameters of the winning model revealed main effects of group (F_1,42_=15.40, p<.001, pFDR<.001), emotion (F_2,84_=653.88, p<.001, pFDR<.001), and movement (F_1,42_=129.80, p<.001, pFDR<.001) as well as significant pairwise interactions between all factors (group x movement: F_2,84_=15.76, p<.001, pFDR<.001; group x emotion: F_2,84_=4.10, p=.019, pFDR=.019; emotion x movement: F_2,84_=134.10, p<.001, pFDR<.001). These effects were nonetheless qualified by a significant emotion x movement x group interaction (F_2,84_=11.31, p<.001, pFDR<.001), paralleling reaction times.

We subsequently tested which parameters differed between groups (Table 3). Controls had negative response drifts when pulling angry faces close, whereas the mean value for this parameter was centered around zero in VMPFC patients (Fig. 3B, left; t_17.46_=3.51, p=.002). VMPFC patients also showed lower drift rates than control participants when pushing angry faces away (Fig. 3B, right; t_15.56_=2.92, p=.010). Therefore, response drifts in VMPFC patients were weaker when avoiding angry faces and relatively less negative (i.e. centered around null) when approaching them. VMPFC patients also had reduced response drifts when pushing happy faces away (Fig. 2C, right; t_41.99_=2.29, p=.026), but not when pulling them close (Fig. 2C, left; p=.258). This pattern was also present at trend level for neutral expressions (Fig. 2D; avoid: t_39.10_=1.85, p=.070; approach: p=.723). Thus, VMPFC patients had generally lower drift rates than controls during avoidance movements, especially for angry faces.

**Figure 3.**
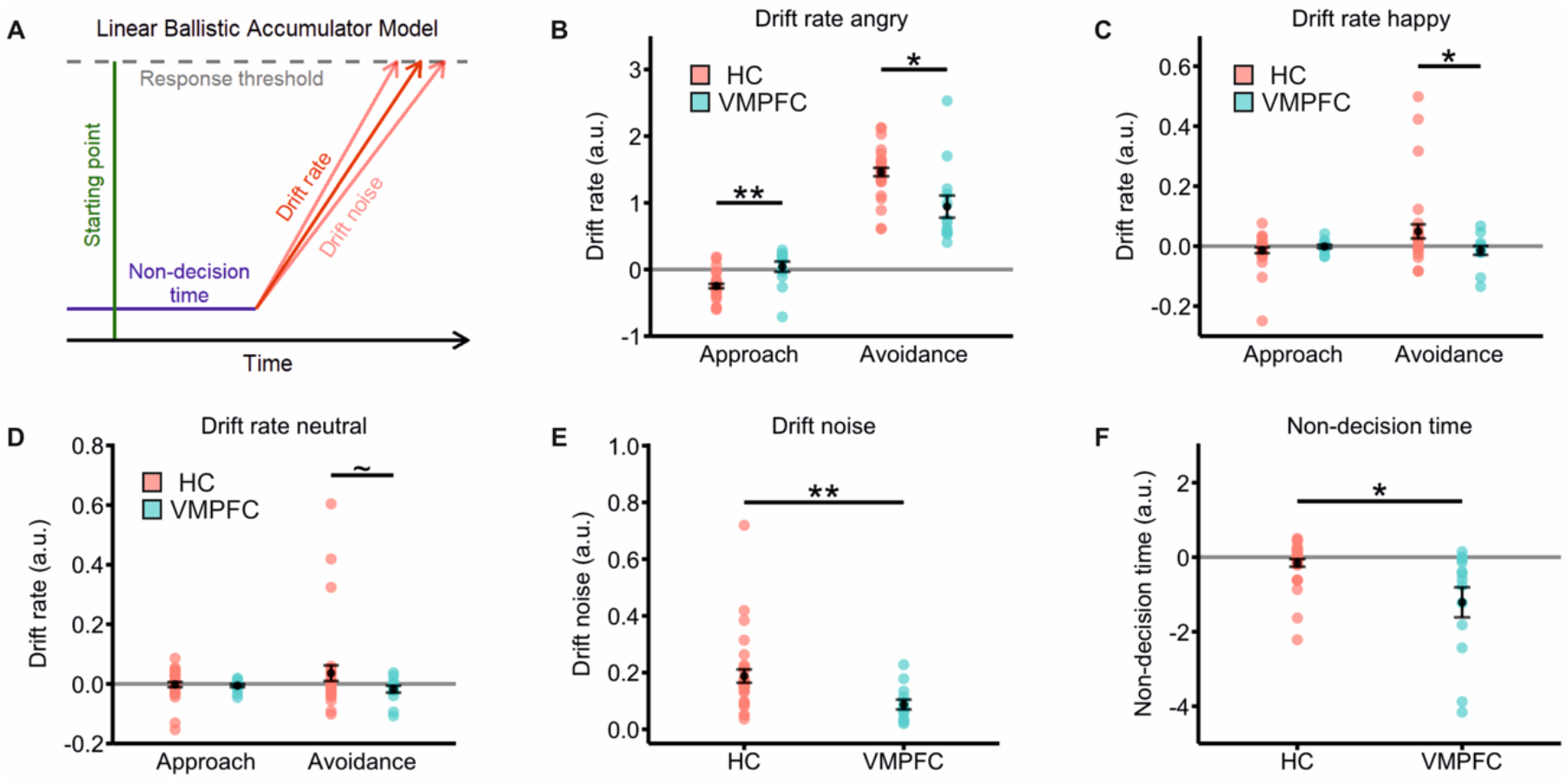
**A)** Schematic depiction of a Linear Ballistic Accumulator (LBA) model, which operationalizes decisions as the result of a sequential evidence accumulation process. The model assumes separate, competing accumulators for each response option, with faster decisions when the response threshold is lower, starting point is higher, non-decision time is shorter, and the drift towards a given option is stronger and less variable (i.e. higher drift rate and lower drift noise). We estimated the parameters from each participant’s reaction time distribution with a maximum likelihood algorithm. **B)** Ventromedial prefrontal cortex (VMPFC) patients showed less negative (i.e. around zero) drift rates than healthy controls (HC) when pulling angry faces close (left), and lower drift rates when pushing angry faces away (right). **C)** VMPFC patients had lower drift rates than HC when pushing happy faces away. **D)** VMPFC patients displayed trend-level lower drift rates than HC when avoiding neutral faces. **E)** VMPFC patients had lower drift noise. **F)** VMPFC patients showed shorter non-decision times. A.u.: arbitrary units. ~p<.1, *p<.05, **p<.01.

**Table 3:** Group-wise means and standard errors of free parameters from the winning model

| Parameter | HC | VMPFC |
| --- | --- | --- |
| Drift rate pull angry** | $-.24 \pm .03$ | $.04 \pm .07$ |
| Drift rate pull happy | $-.01 \pm .009$ | $-.001 \pm .005$ |
| Drift rate pull neutral | $-.002 \pm .008$ | $-.005 \pm .005$ |
| Drift rate push angry* | $1.46 \pm .06$ | $.94 \pm .16$ |
| Drift rate push happy* | $.04 \pm .02$ | $-.01 \pm .01$ |
| Drift rate push neutral | $.03 \pm .02$ | $-.01 \pm .01$ |
| Drift noise** | $.18 \pm .02$ | $.08 \pm .01$ |
| Starting point | $.13 \pm .02$ | $.08 \pm .03$ |
| Threshold | $.87 \pm .14$ | $1.56 \pm .39$ |
| Non-decision time* | $-.15 \pm .10$ | $-1.21 \pm .40$ |

HC: healthy controls; VMPFC: orbitofrontal cortex patients. Asterisks denote significant between-group differences in parameter estimates at \*p<.05 or \*\*p<.01.

Regarding the remaining parameters, the patient group displayed reduced drift noise (Fig. 2E; t_41.11_= 3.42, p=.001; HC: 0.18± 0.02, VMPFC: 0.08±0.01), and non-decision times (Fig. 2F; t_13.58_=2.54, p=.023; HC: −.15±.10, VMPFC: −1.21±.40). There were no group differences in decision threshold (p=.126) or starting point (p=.364). Hence, evidence accumulation began earlier and was less variable across conditions in VMPFC patients.

In a final exploratory analysis, we tested for linear associations between LBA parameters altered in VMPFC patients and clinical impairment. For drift rates, we limited these analyses to threat approach (pull angry minus pull neutral) and threat avoidance (push angry minus push happy), as these were the same contrasts that we computed for correlations with reaction times. We also correlated drift noise and non-decision times with clinical scores. There were no associations between either score and any of the clinical scales (all p>.234).

## Discussion

Maladaptive social behavior is common after ventromedial prefrontal cortex (VMPFC) damage (Anderson et al., 2006; Blair, 2004), but the neurocognitive processes underlying these symptoms remain elusive. Here, we tested whether patients with acquired VMPFC lesions show altered automatic responses to emotional facial expressions. VMPFC patients displayed both reduced avoidance of, and increased approach to angry faces. Modelling of reaction times indicated that these biases are due to differences in stimulus processing rather than to pre-existing preferences for either type of stimuli. Between-group comparisons revealed relatively slower evidence accumulation when avoiding angry faces in VMPFC patients relative to controls. Moreover, patients lacked the negative response drifts that controls showed during approach of angry expressions. VMPFC patients further evidenced less variable and earlier-starting evidence accumulation. The approach bias in VMPFC patients was associated with self-reported clinical measures of impulsive and disinhibited behavior. Patients also committed more errors, which was in turn correlated with greater self-reported impulsivity, disinhibition, problems in emotional control, and worse self-monitoring. Finally, larger lesions were linked with a relatively more pronounced approach bias to angry faces, but not with error rates or avoidance biases. All in all, these findings suggest that VMPFC damage can precipitate maladaptive behavior by altering the implicit processing of threatening social information during action selection.

### VMPFC lesions increase approach and reduce avoidance of threatening stimuli

Our findings expand on a previous report indicating that VMPFC-damaged individuals report negative facial expressions to be more approachable (Willis et al., 2010). Here, we showed that this translates into observable, automatic motor behavior, such that VMPFC patients were quicker to actively approach angry faces (i.e., pull them towards themselves), but slower to avoid them (i.e., push them away). Reduced implicit avoidance of angry faces has been reported in psychopathic offenders (von Borries et al., 2012), who also display dampened physiological reactivity to threatening distractors (Newman et al., 2010) and deficits in VMPFC-dependent tasks (Blair, 2010). Therefore, VMPFC dysfunction seems to confer both lower threat aversion and enhanced threat approach, features that may facilitate antisocial behavior in some individuals.

The present results broadly converge with clinical (Blair, 2004), volumetric (Chester et al., 2017), and functional (Beyer et al., 2015; Gilam et al., 2015) studies asserting that the VMPFC is essential for the regulation of aggressive urges. Our data further indicate that the VMPFC does not merely suppress automatic impulses but rather directs the course of approach-avoidance reactions, in line with recent proposals (Hiser & Koenigs, 2018; Rudebeck & Rich, 2018), and with the well-known association between damage to this region and disadvantageous decision-making (Koenigs & Tranel, 2007). Given that the VMPFC is involved in the anticipation and evaluation of actions related to certain stimuli (R. C. Wilson et al., 2014), we suggest that VMPFC dysfunction gives rise to an altered processing of threat signals. Specifically, it might be that VMPFC damage compromises the prediction of behavioral outcomes associated with potentially punishing stimuli, i.e., tagging angry faces as neutral or even potentially rewarding (Rudebeck & Murray, 2014). These abnormal value forecasts can in turn enable the impulsive, rule-breaking behavior that characterizes the sequelae of some VMPFC lesions.

In line with the latter statement, approach towards angry relative to neutral faces was linked with greater self-reported disinhibition and impulsive behavior. Paralleling our results, it has been reported that patients with borderline personality disorder, who regularly engage in antagonistic and aggressive behavior, also show an approach bias to angry faces (Bertsch et al., 2018) and comparable levels of impulsivity and self-reported anger as those of VMPFC patients (Berlin et al., 2005). Similarly, healthy individuals with high trait anger are quicker to approach angry relative to happy faces (Veenstra et al., 2017). The current results thus provide further evidence that threat signals might act as appetitive stimuli for individuals with externalizing symptomatology (Chester, 2017), and further add that VMPFC lesions might precipitate such dysfunctional evaluation processes. Although VMPFC patients’ externalizing behavior is typically less severe than that of clinically antisocial individuals (Berlin et al., 2005), lesions to the VMPFC are the most frequently associated with subsequent criminality (Darby et al., 2018). Therefore, VMPFC lesions are likely to confer at least some risk for antisocial behavior through other mediating features such as problems in value representation (Blair, 2010), as the present data support.

Of note, the response tendencies observed in VMPFC patients were independent of gaze direction. This pattern deviates from previous studies reporting group-specific approach-avoidance biases exclusively for directly-gazing angry faces (Roelofs et al., 2010; von Borries et al., 2012). Hence, the present findings tentatively suggest that VMPFC lesions might be associated with reduced sensitivity to gaze direction. We did find, however, that straight-looking neutral faces were linked with slower reaction times across the whole sample irrespective of movement type. The latter observation insinuates that neutral expressions, due to their inherent ambiguity (Blasi et al., 2009), are more thoroughly evaluated when directed to oneself.

Importantly, VMPFC patients performed generally worse than controls in the implicit approach-avoidance task (AAT), which is suggestive of difficulties in ignoring task-irrelevant stimulus features. This observation concurs with other studies in showing that VMPFC patients are more susceptible to distraction by to-be-ignored stimulus characteristics (Kuusinen et al., 2018; Mäki-Marttunen et al., 2017), and agrees with the general idea that VMPFC damage hinders the implementation of goal-directed behavior (Rudebeck & Rich, 2018). Moreover, error rates were associated with greater self-reported impulsivity and disinhibition in VMPFC patients, as well as with worse emotional control and self-monitoring. Such findings speak for the predictive validity of the AAT and support its potential usefulness for assessing emotional dysfunction in neurological patients (Fricke & Vogel, 2020).

Neuroanatomical analyses showed that lesion size strongly predicted threat approach, and that no specific prefrontal regions accounted for this effect. These analyses confirmed nevertheless that lesions were mostly localized in the VMPFC, with additional extensive damage to the frontal pole. Thus, the current data suggest that ventromedial and anterior prefrontal cortex are most strongly linked with the observed threat approach bias. It has been suggested that all prefrontal subregions carry out evaluative functions, with a relative local specialization for certain types of information (Dixon et al., 2017). In the context of emotional control, the frontal pole has been postulated to monitor current and alternative strategies in order to facilitate response switching as dictated by current task demands, e.g. from approach to avoidance (Bramson et al., 2020; Koch et al., 2018). Therefore, we speculate that the implementation of adaptive approach-avoidance behavior might rely on both VMPFC-dependent value representations as well as emotional action monitoring in frontopolar areas.

### VMPFC lesions affect latent decision parameters

We used Linear Ballistic Accumulator (LBA) modelling to delve deeper into the decision processes underlying approach-avoidance responses in VMPFC patients. These analyses indicated that emotional facial expressions modulated drift rates (i.e., the speed of evidence accumulation after a stimulus appears) but no other parameters. These findings extend previous drift diffusion modelling work using an explicit version of the AAT in which emotional expressions impacted not only drift rates but also response thresholds and non-decision times (Tipples, 2019). Hence, the influence of emotional expressions on latent decision variables may be less pronounced when facial expressions are to be ignored. The present data do however fully dovetail previous modelling studies in that response drifts were maximal when threatening stimuli were to be avoided (Krypotos et al., 2015; Mennella et al., 2020; Tipples, 2019). Our results complement these findings by showing that angry faces automatically bias evidence accumulation towards avoidance even in the absence of explicit response contingencies.

Between-group comparisons of model parameters revealed profound differences between VMPFC patients and control participants. VMPFC patients showed near-zero drift rates when approaching (i.e., pulling) angry faces, whereas healthy controls showed negative values in this parameter. VMPFC lesions might thus eliminate a default bias against threat approach. In addition, we observed weaker response drifts during avoidance responses (i.e., push movements) in patients relative to controls. The group difference in this parameter was strongest for angry facial expressions but also present in happy and, at trend level, neutral trials. Evidence accumulation leading to avoidance decisions is hence more sluggish in VMPFC patients, and especially so in the presence of angry facial expressions. Therefore, the incongruent approach behavior often observed in VMPFC patients (Perry et al., 2016; Willis et al., 2010) might be partly attributable to an altered evidence accumulation process in response to social signals. Specifically, evidence accumulation in VMPFC patients seems to lack a bias against threat approach and is generally slower during avoidance. In control participants, in contrast, the positive drift rates when pushing angry faces away might have outweighed the negative drifts when pulling them close, resulting in threat avoidance. These observations agree with the idea that the VMPFC encodes the currently relevant state-space (Stalnaker et al., 2015; R. C. Wilson et al., 2014). Angry facial expressions should, on the basis of previous experience, evoke a representation of possible negative outcomes and thereby facilitate avoidance, as seen in the reaction times of control participants. This negative outcome representation is abolished after VMPFC lesions, presumably producing the observed alterations in evidence accumulation and the resulting abnormal approach-avoidance responses.

In addition, VMPFC patients displayed relatively shorter non-decision times and lower drift rate variability irrespective of experimental condition. This implies that approach-avoidance decision processes start earlier and are more rigid in VMPFC patients as compared to control participants. The lower non-decision times are in consonance with the generally speeded responding and higher error rates incurred by VMPFC patients, as well as with the enhanced impulsivity often observed in VMPFC-damaged individuals (Berlin et al., 2004, 2005). On the other hand, the reduced drift rate variability observed in patients parallels the deficits in goal-directed behavior subsequent to VMPFC damage, i.e., a failure to update stimulus value resulting in perseverative responses (Rudebeck et al., 2013; Rudebeck & Murray, 2014).

Surprisingly, clinical impairment could be predicted by reaction times but not by any single LBA parameter. Thus, VMPFC patients’ emotional dysregulation appears to be more strongly influenced by the combination of multiple latent decision processes (which putatively produce the observed reaction times) rather than by any single one of them in isolation. The absence of group differences in starting point or decision threshold is also noteworthy, as it indicates that the approach bias observed in VMPFC patients is likely due to post-stimulus processing rather than to pre-existing response tendencies. This finding agrees with a recent modeling study on approach-avoidance decisions in which the emotional valence of the stimuli modulated drift rates but no other parameters in the model (Mennella et al., 2020). Taken together, LBA results suggest that damage to the VMPFC might lead to rapid and invariant evidence accumulation, which is in turn slower when avoiding threatening stimuli but relatively faster when approaching these signals.

### Limitations

The cross-sectional nature of the design, along with the reduced sample size common in studies with focal lesion patients (Motzkin et al., 2015; Pujara et al., 2016), constrain the generalizability of the present results. Special caution should be exercised regarding the correlations: even though we used bootstrapping to assess their robustness, the ability of the implicit AAT to track interindividual differences is uncertain due to the lack of data on this instrument’s reliability (Hedge et al., 2018). In general, effect sizes from discovery studies such as the present one should be assumed to be inflated until replication or follow up studies permit a more precise estimation of the true effect (B. M. Wilson et al., 2020). It should also be noted that there was no lesion control group, which renders the regional specificity of the results uncertain. Nonetheless, anatomical analyses revealed that the strongest lesion overlap was located in ventromedial and rostral-anterior aspects. Finally, due to time constraints, we were not able to measure patients’ explicit emotion recognition abilities, which are sometimes (Heberlein et al., 2008) but not always (Willis et al., 2010) impaired in VMPFC patients. Control tasks involving emotion recognition and distraction susceptibility as well as additional measures such as eye-tracking (Goursaud & Bachevalier, 2020) would permit a more precise dissection of the mechanisms underlying altered approach-avoidance behavior after VMPFC damage. This limitation is minimized by the fact that the task did not require emotion recognition to be performed.

### Conclusion

The present study provides insight on how VMPFC dysfunction impacts the processing of threatening information during approach-avoidance decisions. This was manifested in altered evidence accumulation in response to threatening stimuli in combination with markers of premature and inflexible decision-making. Intervention programs to improve social functioning in VMPFC patients might therefore benefit from a focus on correctly interpreting and reacting to emotional information as well as on ameliorating impulsivity (Levine et al., 2008). In sum, our study demonstrates that VMPFC damage can steer individuals towards maladaptive approach behavior by biasing the automatic evaluation of threat signals.

## Acknowledgements and funding

This study was funded by the German Science Foundation (grant number KR3691/5-1). MBR was supported by a German Science Foundation stipend (grant number BU 3756/1-1). AKS, TE, and IF were supported by the Research Council of Norway through a grant (project number 240389) and through the Centres of Excellence scheme (project number 262762 RITMO). KR was supported by a VICI grant (#453-12-001) from the Netherlands Organization for Scientific Research (NWO) and a consolidator grant from the European Research Council (ERC_CoG-2017_772337). We thank Dr. Torstein R. Meling for help with patient recruitment and Dr. Per Kristian Hol for clinical evaluation of the patients’ MRI scans. Dr. Buades-Rotger, Dr. Solbakk, Dr. Liebrand, Dr. Endestad, Dr. Funderud, Mr. Siegwardt, Ms. Enter, Dr. Roelofs, and Dr. Krämer report no biomedical financial interests or potential conflicts of interest.

## Notes

### Competing Interest Statement

The authors have declared no competing interest.

### Summary of Updates

We implemented substantial changes after peer review and subsequent acceptance at the Journal of Cognitive Neuroscience, most notably: - We performed new analyses of lesion anatomy, adding a new table. - We refined the regional specificity of our claims on the basis of the new results, and updated figures accordingly. - We ran an additional confirmatory analysis of variance on model parameters.

## References

Anderson, S. W., Barrash, J., Bechara, A., & Tranel, D. (2006). Impairments of emotion and real-world complex behavior following childhood- or adult-onset damage to ventromedial prefrontal cortex. Journal of the International Neuropsychological Society, 12(2), 224–235. https://doi.org/10.1017/S1355617706060346

Barrash, J., Tranel, D., & Anderson, S. W. (2000). Acquired Personality Disturbances Associated With Bilateral Damage to the Ventromedial Prefrontal Region. Developmental Neuropsychology, 18(3), 355–381. https://doi.org/10.1207/S1532694205Barrash

Beer, J. S., John, O. P., Scabini, D., & Knight, R. T. (2006). Orbitofrontal cortex and social behavior: Integrating self-monitoring and emotion-cognition interactions. Journal of Cognitive Neuroscience, 18(6), 871–879. PubMed. https://doi.org/10.1162/jocn.2006.18.6.871

Berlin, H. A., Rolls, E. T., & Iversen, S. D. (2005). Borderline Personality Disorder, Impulsivity, and the Orbitofrontal Cortex. American Journal of Psychiatry, 162(12), 2360–2373. https://doi.org/10.1176/appi.ajp.162.12.2360

Berlin, H. A., Rolls, E. T., & Kischka, U. (2004). Impulsivity, time perception, emotion and reinforcement sensitivity in patients with orbitofrontal cortex lesions. Brain, 127(5), 1108–1126. https://doi.org/10.1093/brain/awh135

Bertsch, K., Roelofs, K., Roch, P. J., Ma, B., Hensel, S., Herpertz, S. C., & Volman, I. (2018). Neural correlates of emotional action control in anger-prone women with borderline personality disorder. Journal of Psychiatry & Neuroscience: JPN, 43(3), 161–170. Pmc. https://doi.org/10.1503/jpn.170102

Beyer, F., Münte, T. F., Göttlich, M., & Krämer, U. M. (2015). Orbitofrontal Cortex Reactivity to Angry Facial Expression in a Social Interaction Correlates with Aggressive Behavior. Cerebral Cortex, 25(9), 3057–3063. https://doi.org/10.1093/cercor/bhu101

Blair, R. J. R. (2004). The roles of orbital frontal cortex in the modulation of antisocial behavior. Brain and Cognition, 55(1), 198–208. https://doi.org/10.1016/S0278-2626(03)00276-8

Blair, R. J. R. (2010). Psychopathy, frustration, and reactive aggression: The role of ventromedial prefrontal cortex. British Journal of Psychology (London, England: 1953), 101(Pt 3), 383–399. https://doi.org/10.1348/000712609X418480

Blasi, G., Hariri, A. R., Alce, G., Taurisano, P., Sambataro, F., Das, S., Bertolino, A., Weinberger, D. R., & Mattay, V. S. (2009). Preferential amygdala reactivity to the negative assessment of neutral faces. Biological Psychiatry, 66(9), 847–853. PubMed. https://doi.org/10.1016/j.biopsych.2009.06.017

Bramson, B., Folloni, D., Verhagen, L., Hartogsveld, B., Mars, R. B., Toni, I., & Roelofs, K. (2020). Human Lateral Frontal Pole Contributes to Control over Emotional Approach–Avoidance Actions. Journal of Neuroscience, 40(14), 2925–2934. https://doi.org/10.1523/JNEUROSCI.2048-19.2020

Brown, S. D., & Heathcote, A. (2008). The simplest complete model of choice response time: Linear ballistic accumulation. Cognitive Psychology, 57(3), 153–178. PubMed. https://doi.org/10.1016/j.cogpsych.2007.12.002

Burnham, K. P., & Anderson, D. R. (2004). Multimodel Inference: Understanding AIC and BIC in Model Selection. Sociological Methods & Research, 33(2), 261–304. https://doi.org/10.1177/0049124104268644

Chester, D. S. (2017). The Role of Positive Affect in Aggression. Current Directions in Psychological Science, 26(4), 366–370. https://doi.org/10.1177/0963721417700457

Chester, D. S., Lynam, D. R., Milich, R., & DeWall, C. N. (2017). Physical aggressiveness and gray matter deficits in ventromedial prefrontal cortex. Cortex, 97, 17–22. https://doi.org/10.1016/j.cortex.2017.09.024

Clithero, J. A., & Rangel, A. (2014). Informatic parcellation of the network involved in the computation of subjective value. Social Cognitive and Affective Neuroscience, 9(9), 1289–1302. https://doi.org/10.1093/scan/nst106

Cramer, A. O. J., van Ravenzwaaij, D., Matzke, D., Steingroever, H., Wetzels, R., Grasman, R. P. P. P., Waldorp, L. J., & Wagenmakers, E.-J. (2016). Hidden multiplicity in exploratory multiway ANOVA: Prevalence and remedies. Psychonomic Bulletin & Review, 23(2), 640–647. https://doi.org/10.3758/s13423-015-0913-5

Darby, R. R., Horn, A., Cushman, F., & Fox, M. D. (2018). Lesion network localization of criminal behavior. Proceedings of the National Academy of Sciences, 115(3), 601. https://doi.org/10.1073/pnas.1706587115

Davidson, R. J., Putnam, K. M., & Larson, C. L. (2000). Dysfunction in the Neural Circuitry of Emotion Regulation—A Possible Prelude to Violence. Science, 289(5479), 591–594. https://doi.org/10.1126/science.289.5479.591

Dixon, M. L., Thiruchselvam, R., Todd, R., & Christoff, K. (2017). Emotion and the prefrontal cortex: An integrative review. Psychological Bulletin, 143(10), 1033–1081. https://doi.org/10.1037/bul0000096

Ekman, P., & Friesen, W. V. (1976). Pictures of Facial Affect. Consulting Psychologists Press. https://ci.nii.ac.jp/naid/10011335061/

Folloni, D., Sallet, J., Khrapitchev, A. A., Sibson, N., Verhagen, L., & Mars, R. B. (2019). Dichotomous organization of amygdala/temporal-prefrontal bundles in both humans and monkeys. ELife, 8, e47175. https://doi.org/10.7554/eLife.47175

Fricke, K., & Vogel, S. (2020). How interindividual differences shape approach-avoidance behavior: Relating self-report and diagnostic measures of interindividual differences to behavioral measurements of approach and avoidance. Neuroscience & Biobehavioral Reviews, 111, 30–56. https://doi.org/10.1016/j.neubiorev.2020.01.008

Gilam, G., Lin, T., Raz, G., Azrielant, S., Fruchter, E., Ariely, D., & Hendler, T. (2015). Neural substrates underlying the tendency to accept anger-infused ultimatum offers during dynamic social interactions. NeuroImage, 120(Supplement C), 400–411. https://doi.org/10.1016/j.neuroimage.2015.07.003

Goursaud, A.-P. S., & Bachevalier, J. (2020). Altered face scanning and arousal after orbitofrontal cortex lesion in adult rhesus monkeys. Behavioral Neuroscience, 134(1), 45–58. https://doi.org/10.1037/bne0000342

Gross, J. J. (2015). Emotion Regulation: Current Status and Future Prospects. Psychological Inquiry, 26(1), 1–26. https://doi.org/10.1080/1047840X.2014.940781

Heathcote, A., & Hayes, B. (2012). Diffusion versus linear ballistic accumulation: Different models for response time with different conclusions about psychological mechanisms? Canadian Journal of Experimental Psychology = Revue Canadienne de Psychologie Experimentale, 66(2), 125–136. PubMed. https://doi.org/10.1037/a0028189

Heberlein, A. S., Padon, A. A., Gillihan, S. J., Farah, M. J., & Fellows, L. K. (2008). Ventromedial Frontal Lobe Plays a Critical Role in Facial Emotion Recognition. Journal of Cognitive Neuroscience, 20(4), 721–733. https://doi.org/10.1162/jocn.2008.20049

Hedge, C., Powell, G., & Sumner, P. (2018). The reliability paradox: Why robust cognitive tasks do not produce reliable individual differences. Behavior Research Methods, 50(3), 1166–1186. https://doi.org/10.3758/s13428-017-0935-1

Hiser, J., & Koenigs, M. (2018). The Multifaceted Role of the Ventromedial Prefrontal Cortex in Emotion, Decision Making, Social Cognition, and Psychopathology. Biological Psychiatry, 83(8), 638–647. https://doi.org/10.1016/j.biopsych.2017.10.030

Jenkins, L. M., Andrewes, D. G., Nicholas, C. L., Drummond, K. J., Moffat, B. A., Phal, P. M., & Desmond, P. (2018). Emotional reactivity following surgery to the prefrontal cortex. Journal of Neuropsychology, 12(1), 120–141. https://doi.org/10.1111/jnp.12110

Jenkins, L. M., Andrewes, D. G., Nicholas, C. L., Drummond, K. J., Moffat, B. A., Phal, P. M., Desmond, P., & Kessels, R. P. C. (2014). Social cognition in patients following surgery to the prefrontal cortex. Psychiatry Research: Neuroimaging, 224(3), 192–203. https://doi.org/10.1016/j.pscychresns.2014.08.007

Koch, S. B. J., Mars, R. B., Toni, I., & Roelofs, K. (2018). Emotional control, reappraised. Neuroscience & Biobehavioral Reviews, 95, 528–534. https://doi.org/10.1016/j.neubiorev.2018.11.003

Koenigs, M., & Tranel, D. (2007). Irrational economic decision-making after ventromedial prefrontal damage: Evidence from the Ultimatum Game. The Journal of Neuroscience: The Official Journal of the Society for Neuroscience, 27, 951–956. https://doi.org/10.1523/JNEUROSCI.4606-06.2007

Krypotos, A.-M., Beckers, T., Kindt, M., & Wagenmakers, E.-J. (2015). A Bayesian hierarchical diffusion model decomposition of performance in Approach–Avoidance Tasks. Cognition and Emotion, 29(8), 1424–1444. https://doi.org/10.1080/02699931.2014.985635

Kuusinen, V., Cesnaite, E., Peräkylä, J., Ogawa, K. H., & Hartikainen, K. M. (2018). Orbitofrontal Lesion Alters Brain Dynamics of Emotion-Attention and Emotion-Cognitive Control Interaction in Humans. Frontiers in Human Neuroscience, 12. https://doi.org/10.3389/fnhum.2018.00437

Levine, B., Turner, G. R., & Stuss, D. T. (2008). Rehabilitation of frontal lobe functions. In Cognitive neurorehabilitation: Evidence and application, 2nd ed (pp. 464–486). Cambridge University Press. https://doi.org/10.1017/CBO9781316529898.033

Løvstad, M., Funderud, I., Endestad, T., Due-Tønnessen, P., Meling, T. R., Lindgren, M., Knight, R. T., & Solbakk, A. K. (2012). Executive functions after orbital or lateral prefrontal lesions: Neuropsychological profiles and self-reported executive functions in everyday living. Brain Injury, 26(13–14), 1586–1598. PubMed. https://doi.org/10.3109/02699052.2012.698787

Lucantonio, F., Stalnaker, T. A., Shaham, Y., Niv, Y., & Schoenbaum, G. (2012). The impact of orbitofrontal dysfunction on cocaine addiction. Nature Neuroscience, 15(3), 358–366. https://doi.org/10.1038/nn.3014

Lundqvist, D., Flykt, A., & Öhman, A. (1998). The Karolinska directed emotional faces (KDEF). CD ROM from Department of Clinical Neuroscience, Psychology Section, Karolinska Institutet, 91, 630.

Mäki-Marttunen, V., Kuusinen, V., Peräkylä, J., Ogawa, K. H., Brause, M., Brander, A., & Hartikainen, K. M. (2017). Greater Attention to Task-Relevant Threat Due to Orbitofrontal Lesion. Journal of Neurotrauma, 34(2), 400–413. https://doi.org/10.1089/neu.2015.4390

Maldjian, J. A., Laurienti, P. J., Kraft, R. A., & Burdette, J. H. (2003). An automated method for neuroanatomic and cytoarchitectonic atlas-based interrogation of fMRI data sets. NeuroImage, 19(3), 1233–1239. https://doi.org/10.1016/S1053-8119(03)00169-1

Mennella, R., Vilarem, E., & Grèzes, J. (2020). Rapid approach-avoidance responses to emotional displays reflect value-based decisions: Neural evidence from an EEG study. NeuroImage, 222, 117253. https://doi.org/10.1016/j.neuroimage.2020.117253

Motzkin, J. C., Philippi, C. L., Wolf, R. C., Baskaya, M. K., & Koenigs, M. (2015). Ventromedial Prefrontal Cortex Is Critical for the Regulation of Amygdala Activity in Humans. Biological Psychiatry, 77(3), 276–284. https://doi.org/10.1016/j.biopsych.2014.02.014

Newman, J. P., Curtin, J. J., Bertsch, J. D., & Baskin-Sommers, A. R. (2010). Attention moderates the fearlessness of psychopathic offenders. Biological Psychiatry, 67(1), 66–70. Pmc. https://doi.org/10.1016/j.biopsych.2009.07.035

Perry, A., Lwi, S. J., Verstaen, A., Dewar, C., Levenson, R. W., & Knight, R. T. (2016). The role of the orbitofrontal cortex in regulation of interpersonal space: Evidence from frontal lesion and frontotemporal dementia patients. Social Cognitive and Affective Neuroscience, 11(12), 1894–1901. https://doi.org/10.1093/scan/nsw109

Pujara, M. S., Philippi, C. L., Motzkin, J. C., Baskaya, M. K., & Koenigs, M. (2016). Ventromedial Prefrontal Cortex Damage Is Associated with Decreased Ventral Striatum Volume and Response to Reward. The Journal of Neuroscience, 36(18), 5047. https://doi.org/10.1523/jneurosci.4236-15.2016

Raftery, A. E. (1995). Bayesian Model Selection in Social Research. Sociological Methodology, 25, 111–163. JSTOR. https://doi.org/10.2307/271063

Ripollés, P., Marco-Pallarés, J., de Diego-Balaguer, R., Miró, J., Falip, M., Juncadella, M., Rubio, F., & Rodriguez-Fornells, A. (2012). Analysis of automated methods for spatial normalization of lesioned brains. NeuroImage, 60(2), 1296–1306. https://doi.org/10.1016/j.neuroimage.2012.01.094

Roelofs, K., Putman, P., Schouten, S., Lange, W.-G., Volman, I., & Rinck, M. (2010). Gaze direction differentially affects avoidance tendencies to happy and angry faces in socially anxious individuals. Behaviour Research and Therapy, 48(4), 290–294. https://doi.org/10.1016/j.brat.2009.11.008

Rorden, C., & Brett, M. (2000). Stereotaxic Display of Brain Lesions. Behavioural Neurology, 12(4). https://doi.org/10.1155/2000/421719

Roth, R., Isquith, P., & Gioia, G. (2005). Behavior Rating Inventory of Executive Function—Adult Version (BRIEF-A) (Vol. 20). Psychological Assessment Resources.

Rudebeck, P. H., & Murray, E. A. (2014). The Orbitofrontal Oracle: Cortical Mechanisms for the Prediction and Evaluation of Specific Behavioral Outcomes. Neuron, 84(6), 1143–1156. https://doi.org/10.1016/j.neuron.2014.10.049

Rudebeck, P. H., & Rich, E. L. (2018). Orbitofrontal cortex. Current Biology, 28(18), R1083–R1088. https://doi.org/10.1016/j.cub.2018.07.018

Rudebeck, P. H., Saunders, R. C., Prescott, A. T., Chau, L. S., & Murray, E. A. (2013). Prefrontal mechanisms of behavioral flexibility, emotion regulation and value updating. Nature Neuroscience, 16(8), 1140–1145. https://doi.org/10.1038/nn.3440

Stalnaker, T. A., Cooch, N. K., & Schoenbaum, G. (2015). What the orbitofrontal cortex does not do. Nat Neurosci, 18(5), 620–627. https://doi.org/10.1038/nn.3982

Tipples, J. (2019). Recognising and reacting to angry and happy facial expressions: A diffusion model analysis. Psychological Research, 83(1), 37–47. https://doi.org/10.1007/s00426-018-1092-6

Veenstra, L., Schneider, I. K., Bushman, B. J., & Koole, S. L. (2017). Drawn to danger: Trait anger predicts automatic approach behaviour to angry faces. Cognition and Emotion, 31(4), 765–771. https://doi.org/10.1080/02699931.2016.1150256

von Borries, A. K. L., Volman, I., de Bruijn, E. R. A., Bulten, B. H., Verkes, R. J., & Roelofs, K. (2012). Psychopaths lack the automatic avoidance of social threat: Relation to instrumental aggression. Psychiatry Research, 200(2), 761–766. https://doi.org/10.1016/j.psychres.2012.06.026

Waid-Ebbs, J. K., Wen, P.-S., Heaton, S. C., Donovan, N. J., & Velozo, C. (2012). The item level psychometrics of the behaviour rating inventory of executive function-adult (BRIEF-A) in a TBI sample. Brain Injury, 26(13–14), 1646–1657. https://doi.org/10.3109/02699052.2012.700087

Whiteside, S. P., & Lynam, D. R. (2001). The Five Factor Model and impulsivity: Using a structural model of personality to understand impulsivity. Personality and Individual Differences, 30(4), 669–689. https://doi.org/10.1016/S0191-8869(00)00064-7

Whiteside, S. P., Lynam, D. R., Miller, J. D., & Reynolds, S. K. (2005). Validation of the UPPS impulsive behaviour scale: A four-factor model of impulsivity. European Journal of Personality, 19(7), 559–574. https://doi.org/10.1002/per.556

Willis, M. L., Palermo, R., Burke, D., McGrillen, K., & Miller, L. (2010). Orbitofrontal cortex lesions result in abnormal social judgements to emotional faces. Neuropsychologia, 48(7), 2182–2187. https://doi.org/10.1016/j.neuropsychologia.2010.04.010

Wilson, B. M., Harris, C. R., & Wixted, J. T. (2020). Science is not a signal detection problem. Proceedings of the National Academy of Sciences, 117(11), 5559–5567. https://doi.org/10.1073/pnas.1914237117

Wilson, R. C., Takahashi, Y. K., Schoenbaum, G., & Niv, Y. (2014). Orbitofrontal cortex as a cognitive map of task space. Neuron, 81(2), 267–279. https://doi.org/10.1016/j.neuron.2013.11.005

